# Alpha oscillations and travelling waves: signatures of predictive coding?

**DOI:** 10.1101/464933

**Authors:** Andrea Alamia, Rufin VanRullen

## Abstract

Predictive coding is a key mechanism to understand the computational processes underlying brain functioning: in a hierarchical network, higher layers predict the activity of lower layers, and the unexplained residuals (i.e. prediction errors) are sent through. Because of its iterative nature, we wondered whether predictive coding could be related to brain oscillatory dynamics. First, we show that a simple 2-layers predictive coding model of visual cortex, with physiological communication delays between layers, naturally gives rise to alpha-band rhythms, similar to experimental observations. Then, we demonstrate that a multi-layer version of the same model can explain the occurrence of oscillatory travelling waves across layers, both feedforward (during visual stimulation) and backward (during rest). Remarkably, the predictions of our model are matched by the analysis of two independent EEG datasets, in which we observed oscillatory travelling waves in both directions.

## Introduction

Predictive coding is a popular computational paradigm to model sensory information processing in the brain, and it has been proposed to explain several cognitive and physiological observations (Rao and Ballard, 1999). Could it also explain the emergence of alpha-band oscillations and some of their distinguished features (e.g. travelling waves)? Alpha rhythms (8-12Hz) are the most predominant oscillations in the human brain, even though their functional role remains controversial. On the one hand alpha is generally stronger in the absence of visual inputs, or when visual inputs are actively ignored – hence a proposed inhibitory role for alpha rhythms (Bonnefond and Jensen, 2012; Jensen and Mazaheri, 2010; Kizuk and Mathewson, 2017; Klimesch et al., 1998; Sadaghiani and Kleinschmidt, 2016). On the other hand, alpha has been suggested to contribute to the temporal framing of sensory inputs (Valera et al., 1981; VanRullen, 2016), and it can sometimes be positively correlated with visual inputs. For example, a recent study reported strong and long-lasting (up to ~1 s) alpha-band oscillations in the visual Impulse Response Function (IRF; Vanrullen and MacDonald, 2012). The IRF is computed by cross-correlating occipital EEG signals recorded from human subjects watching a dynamic sequence of random (white noise) luminance values with the corresponding stimulus sequence (fig.1A). It is, therefore, a direct reflection of visual sensory processing. The existence of significant input-output correlations at lags of nearly 1s is surprising, given the typical neural time constants (<50ms) and the short-lived nature of visual-evoked responses (<0.5s). Additionally, a recent study from our group (Lozano-Soldevilla and VanRullen, 2017) showed that alpha IRF oscillations propagate as a travelling wave across the cortex in an occipital-to-frontal direction (fig.2A,C). This finding is in line with other recent intracranial studies about alpha-frequency cortical travelling waves (Bahramisharif et al., 2013; Halgren et al., 2017; Muller et al., 2018; Zhang et al., 2018). Yet, the mechanisms underlying such oscillations remain debated. Could it be possible that a common computational principle, predictive coding, gives rise to alpha-band oscillations and their typical travelling wave dynamics?

**Figure 1.**
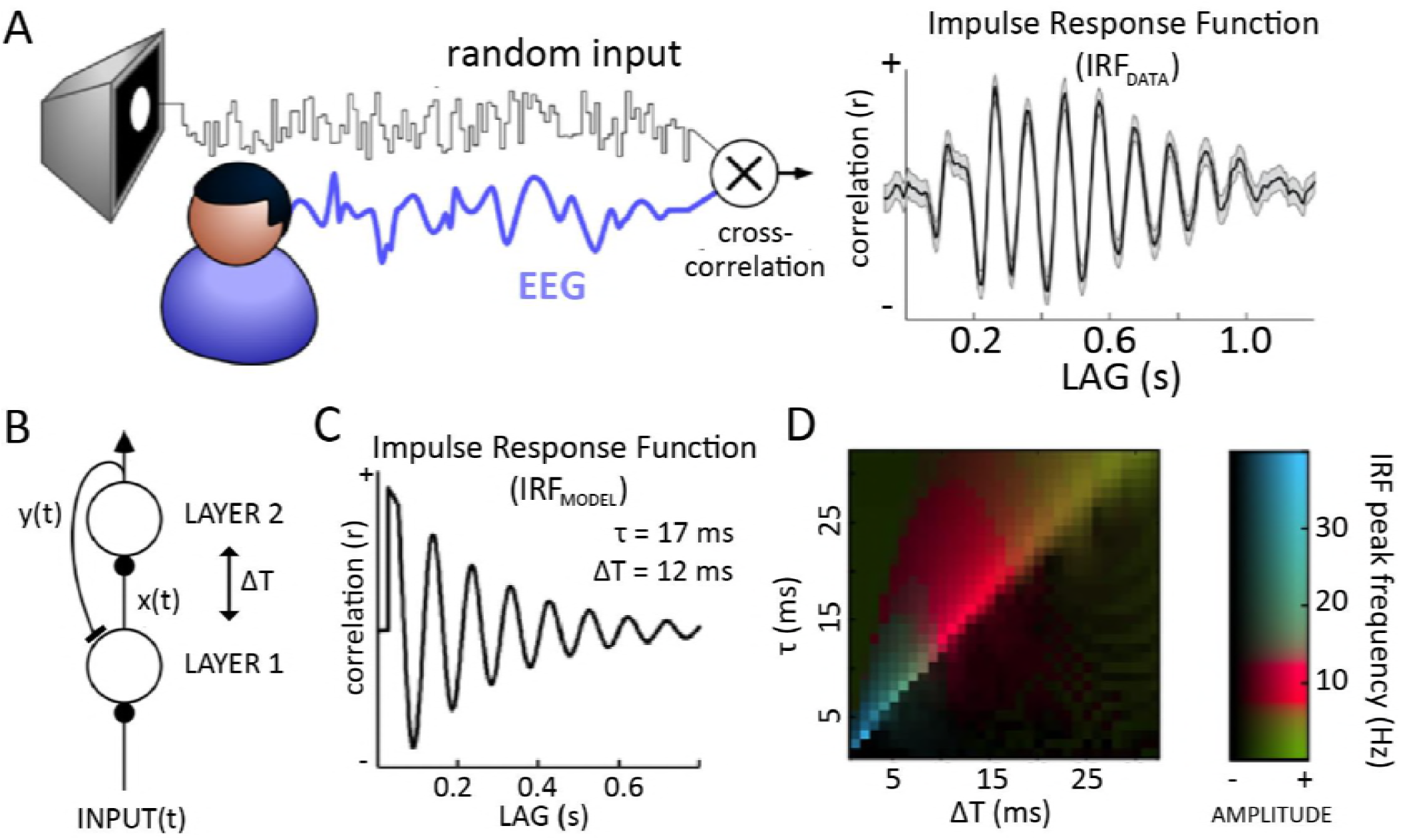
– Alpha-band oscillations in human IRF and in a predictive coding model. A) Cross-correlating the white noise sequence of a stimulus with simultaneously recorded EEG produces an Impulse Response Function (IRF) which reverberates at 10Hz for several successive cycles (one representative subject shown here, electrode POz). B) A simple predictive coding model in which layer 2 makes predictions y(t) about the input received by layer 1, and the residual x(t) (prediction error) is used to update the next prediction. Residual and prediction signals are transmitted to the next or previous layer (respectively) with a communication delay ΔT. Such a model, with physiologically plausible parameters, generates an oscillatory IRF at 10 Hz. C) The oscillatory IRF produced by the model, with communication delay ΔT=12ms, and neural membrane time constant τ=17ms. D) Systematic exploration of these two parameters suggests that alpha reverberation is a robust phenomenon (red colors) within a biologically plausible range of values.

**Figure 2.**
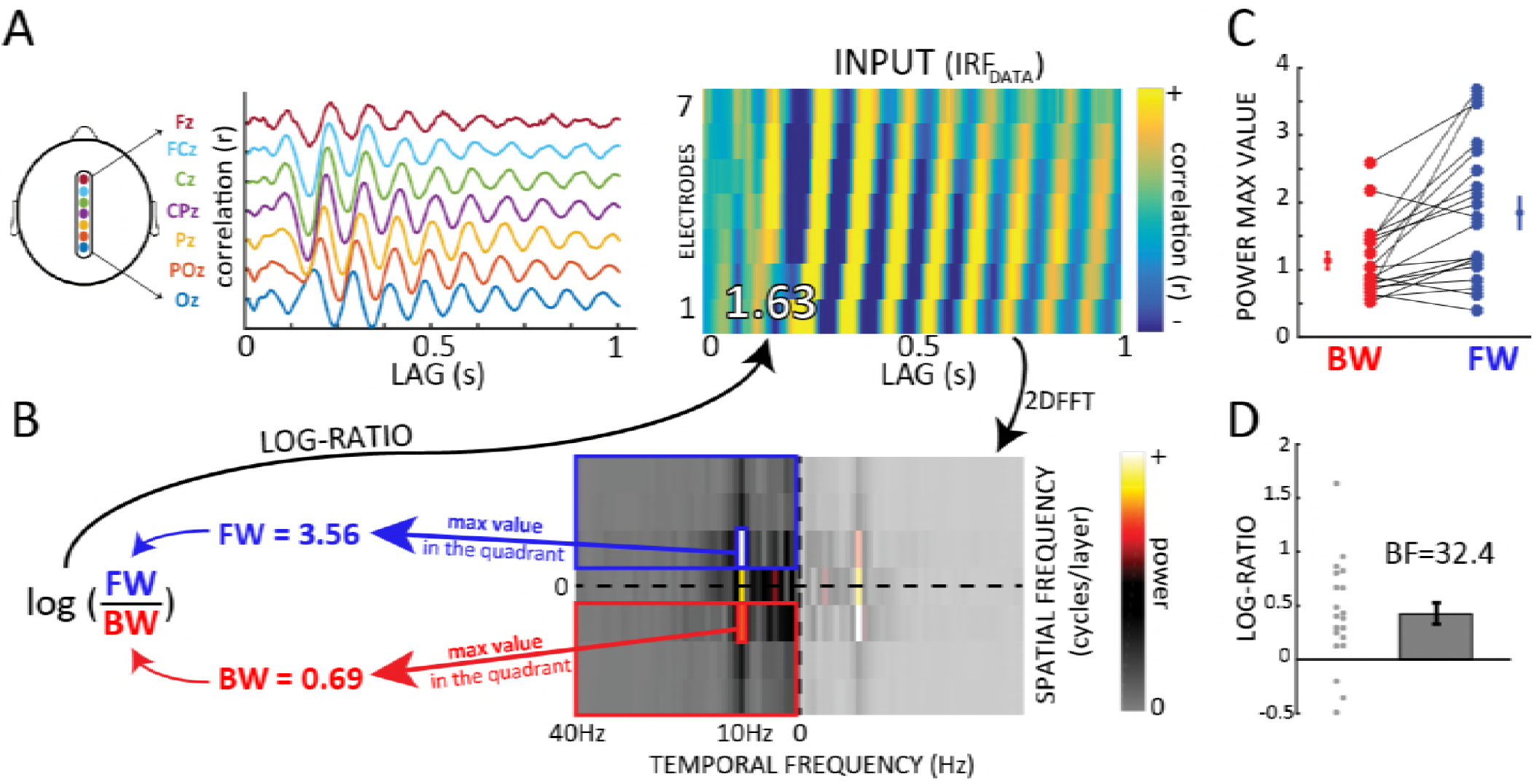
– Alpha-band travelling waves in EEG Impulse Response Functions. A-B) IRF as travelling waves can be observed over the 7 midline electrodes of the 10-20 system (Oz to Fz; one representative subject). From the 2D-map obtained by stacking signals from the 7 electrodes, we compute a 2D-FFT and derive a log-ratio between spectral quadrants that quantifies the direction of the waves. Positive values are associated with feedforward waves (FW), and negative values with backward ones (BW). C) Backward (red) and feedforward (blue) values (max value in the corresponding quadrant of the 2D-FFT) computed over 20 subjects. Participants were watching a white noise luminance sequence (see fig. 1A). D) Log-ratios computed from the values in C. The Bayes Factor (BF) of a one sample t-test against zero confirms the presence of FW waves.

Here, we address this question in two steps. First, we demonstrate the presence of similar long-lasting alpha-band oscillations in the IRF of a simple 2-layers model of visual cortex. Our model does not explicitly integrate or maintain information over extended periods, but merely tries to predict what comes next based on what was just there, i.e., predictive coding. The key insight is that typical neural communication delays between cortical areas can give rise to a reverberation of visual inputs at alpha frequency, as observed experimentally. Second, we expand our predictive coding model by increasing the number of layers to explore the occurrence of alpha band travelling waves in a hierarchical network. After having observed that this hierarchical model produces 1) feedforward (FW) travelling waves during sensory processing and 2) backward (BW) travelling waves in the absence of sensory input, we turned to experimental EEG data to verify these predictions. Remarkably, we found FW waves in human EEG when participants were attending to visual stimulation, and BW waves when participants’ eyes remained closed, in agreement with our model’s predictions.

## Results

### 2-layers model and alpha-band oscillations

In an attempt to explore the emergence of alpha-band IRF oscillations during the visual processing of white-noise luminance sequences (Fig 1A), we initially implemented a simple 2-layers model. The model architecture was inspired by the classic predictive coding model of Rao and Ballard (1999), where each layer attempts to “explain away” (via inhibitory feedback) the activity pattern in the previous layer, which only communicates the “unpredicted” residual signals (via feedforward excitation). Since in our case the stimuli are strictly temporal luminance sequences, without any meaningful spatial arrangement (fig.1A), the original model could be simplified greatly by ignoring spatial selectivity and considering a single neuron (or a single population of neurons) in each of two connected layers (corresponding, e.g. to LGN and V1 of the primate brain). The resulting circuit is illustrated in figure 1B. The specificity of the present approach is to consider the effects of the communication delay ΔT between the two layers (assumed here to be symmetric, for simplicity). The population in layer 1 encodes the residual between the input sequence and the “prediction” received, with a delay ΔT, from layer 2. The instantaneous response *y(t)* of the population in layer 2 is governed by a differential equation (see Materials and Methods) composed of two terms: the first term determines the integration of inputs from layer 1, with a delay ΔT, and the second is a decay term ensuring that neurons eventually return to their resting state in the absence of inputs. The temporal dynamics of neuronal integration and decay are governed by two time constants, respectively τ and τ_decay_, the latter of which was fixed to τ_decay_=200ms. We computed the model’s IRF by cross-correlating each random input luminance sequence *input(t)* with the corresponding layer 2’s output *y(t)*, and averaging the result over 200 trials. That is, we assumed here that layer 2’s activity is an approximation of the EEG signal (however, similar results with only a phase difference were found using layer 1’s activity instead). With biologically plausible values for the τ and ΔT parameters (respectively 17ms and 12ms), the model’s IRF oscillated at a frequency within the alpha-range, thus qualitatively replicating the experimental observations (fig.1C).

### Sensitivity to parameters

We investigated the dependence of IRF oscillations (and of their frequency) on the model’s two free parameters τ and ΔT. First, we employed numerical simulations, similar to the one described above (Fig 1C). A fast-Fourier transform (FFT) applied on the simulated IRF was used to measure the peak oscillatory amplitude and its corresponding frequency. These two measures are color-coded in Figure 1D, displaying the results of a systematic exploration of parameter space. Several combinations of parameters gave rise to oscillatory IRFs (brighter colors); among these, IRF oscillations in the alpha (8-13Hz) frequency range were particularly frequent (red colors). In particular, alpha-band IRF oscillations systematically arose when τ and ΔT lay in their “biologically plausible” range of respectively 15-25ms (Koch et al., 1996; Miyoshi et al., 2010; Rose, 2005; Zaitsev et al., 2012) and 10-15ms (Bair et al., 2002; Nowak et al., 1997; Shimegi et al., 2014).

Second, we also derived an analytical solution to a simplified version of our differential equation system (see Material and Methods). The solution revealed that 1) IRF oscillations occur mainly when the parameter τ lies slightly above ΔT (the optimal oscillatory situation corresponding to τ ≈ 1.27 *ΔT*) and 2) when oscillations occur, the oscillation period equals 8 ∗ ΔT. In other words, alpha-range oscillations (8-13Hz) correspond to ΔT values between 10 and 15ms, and τ values around 15-20ms. Therefore, both of these analytical conclusions directly confirm the results of the numerical simulations.

### Multi-layers model and travelling waves

#### FW travelling waves during visual processing

After having explored the emergence of alpha oscillations in a simple predictive coding model, we investigated whether our model could reproduce other features of alpha-band oscillations, such as their travelling wave dynamics. Consequently, we extended the model to a multi-layers version (fig.3A), in which τ_D_ (decay time constant) remained fixed at 200ms, and ΔT and τ were respectively 12ms and 20ms in all simulations. We chose these values based on the results of the previous parameter space exploration, considering that these parameters are both biologically plausible, and produced IRF alpha oscillations in a 2-layers version of the predictive coding model. As previously, the prediction signals from each layer in our model were treated as the equivalent of EEG signals from distinct electrodes over the human brain, and we used 7 layers to facilitate comparison with experimental data (fig.2A). However, qualitatively similar results were obtained using lower or higher electrode numbers. We created 2D maps by stacking the EEG or IRF signals (x-axis) from the 7 layers (y-axis, see fig. 2B). To quantify the presence and direction of waves, we then extracted the maximum values in the upper and lower quadrants of the 2D-FFT of these maps, representing respectively the amount of feedforward (FW) and backward (BW) signal propagation across layers (fig. 2 A-B). Finally, the log-ratio of these two values quantified the overall direction of the waves: positive ratios indicate predominantly FW waves, whereas negative ratios reveal mostly BW waves.

**Figure 3.**
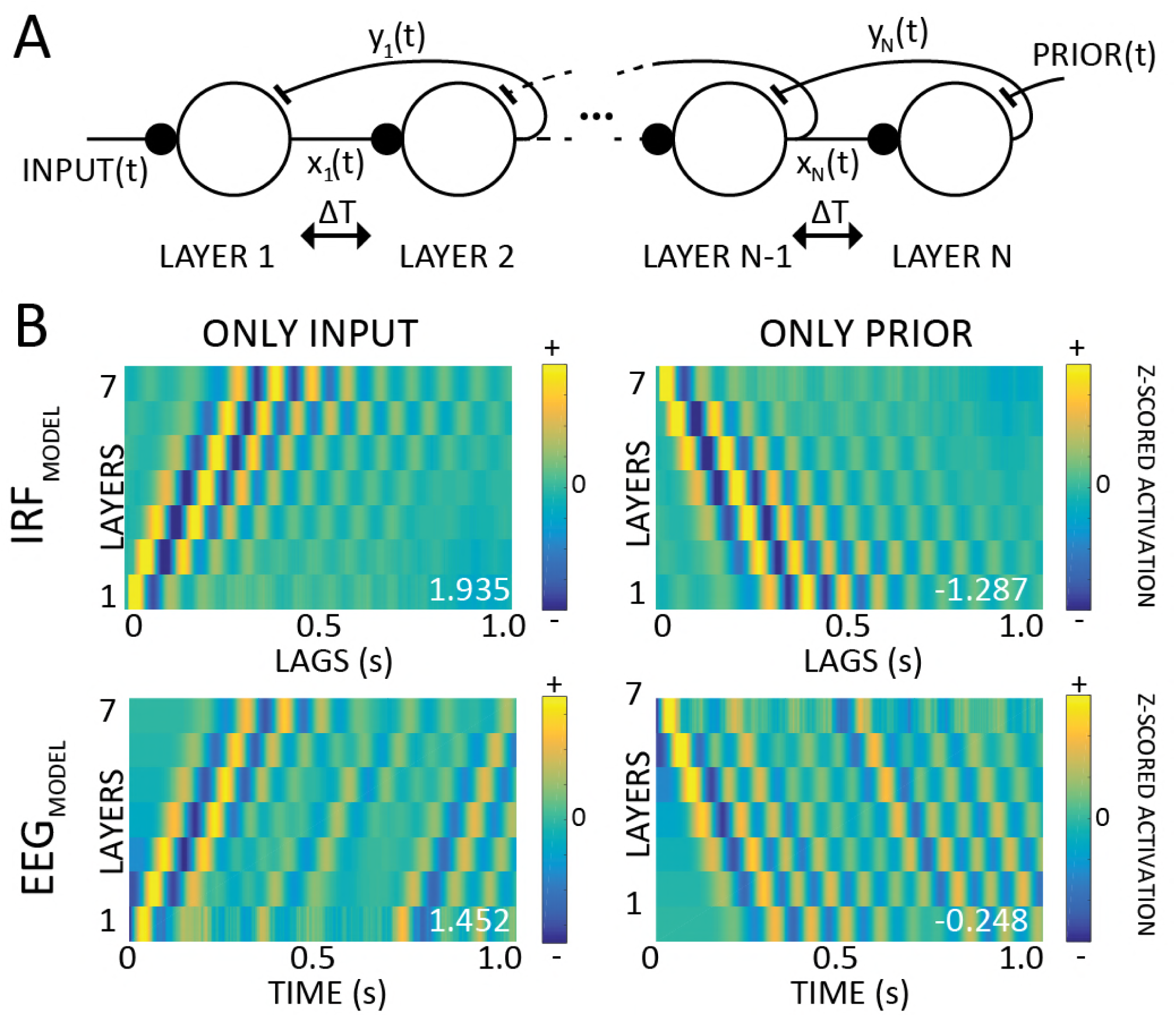
– Alpha-band traveling waves in a hierarchical predictive coding model. A) Multi-layer version of the model: the same parameters ΔT=12ms and τ=20ms are used throughout. The model is fed either a time-varying input (left) or a time-varying prior signal (right), reflecting top-down expectations computed in other parts of the brain. B) 2D maps with only input (left column) or prior signals (right column): travelling waves are visible both in the IRF (computed by cross-correlating prediction signals with input or prior signals, top row) and in the raw prediction signals (considered as a proxy for the EEG, lower row). All the values have been z-scored layer-wise for visualization purposes only. The numbers in white show the log-ratio for each simulation: positive or negative numbers reveal respectively FW or BW waves.

In a first simulation, comparable to the experimental setup described in Figure 1A, we presented the multi-layer model with white-noise inputs only. As shown in the left column of figure 3B, the model response to these inputs displayed feedforward alpha-band oscillatory travelling waves, proceeding from lower to higher layers; this oscillatory propagation was visible both when we applied our wave quantification method to IRF signals (after cross-correlation with the stimulus sequence), and to the raw EEG signals. Whereas on the one hand the presence of FW waves in the model’s IRF (figure 3B, top left panel) was the anticipated model outcome (see Lozano-Soldevilla and VanRullen, 2017 but also fig.2A), on the other hand the presence of travelling waves in the raw (simulated) EEG signal was not readily predictable. Nonetheless, we observed a clear FW propagation in the raw EEG signals of our model, as shown in the bottom-left panel of figure 3B, suggesting that oscillatory travelling waves during sensory processing could be directly visible in EEG recordings (without stimulus cross-correlation). This prediction will be verified in one of the following sections.

#### BW waves in the absence of inputs

Next, we further expanded the model functionality and introduced an endogenous signal at the last layer, namely the prior *y*_*N*+*1(t)*_ (fig.3A – where N is the number of layers). This signal is functionally equivalent to the prediction *y_L_(t)* generated by the model at all previous layers 1 to N, but it is assumed here to arise from higher-level brain regions that are not part of the model, and thus to influence the activity of the last layer as a top-down prediction. In order to investigate the consequences of this new ‘prior’ signal, we first set the model’s input to zero and the prior as a time-varying white-noise signal, allowing us to compute a prior-driven IRF at each level. Note that these design choices were only made to facilitate computations and comparison of input-driven and prior-driven IRFs, and do not reflect any assumption about the statistical structure of inputs or priors under natural conditions of stimulation (natural inputs are not random, and typically do not display a flat power spectrum; similarly, top-down brain signals are unlikely to have white-noise properties). Contrary to the waves’ direction in the previous simulation, the activity of the prior in the absence of inputs generated an oscillatory backward alpha-band travelling wave, which propagated from higher to lower electrodes (figure 3B, top right panel), As previously, we also investigated whether the raw simulated EEG signals would exhibit similar dynamics: as shown in the lower-right panel of figure 3B, we found that this was indeed the case, observing EEG travelling waves proceeding from higher to lower layers. All in all, these results indicate that –at least in the model-EEG recordings are sufficient to identify travelling waves propagating in both directions.

In a final simulation, the model was fed with both the input and the prior, as two independent white-noise signals. In this case, the dynamics induced by the inputs appeared to dominate, which was mathematically expected given τ_D_ >> τ. Consequently, the model revealed predominantly FW waves in this simulation (see fig.S1 for an overview of all possible scenarios).

All in all, the chief conclusion of these simulations is that hierarchical predictive coding gives rise to oscillatory travelling waves that can be seen in both the EEG and IRF signals. Sensory inputs generate FW waves, whereas top-down priors induce BW waves. Although the presence of alpha-frequency FW waves in the IRF during visual processing had been previously demonstrated (Lozano-Soldevilla and VanRullen, 2017; see also fig.2A,C), our model makes further predictions that have not been experimentally validated yet. In the following section, we tested whether these new predictions are matched by experimental EEG data.

### Travelling waves in human EEG data

Our simulations show that hierarchical predictive coding can produce oscillatory travelling waves propagating both feedforward and backward. The model predicts that input processing produces mostly feedforward waves, whereas backward waves are generated by higher-level endogenous signals, and most visible in the absence of visual inputs. Finally, the model suggests that even though cross-correlation and IRFs may be helpful to reveal the travelling waves, their dynamics could also be directly visible in the raw EEG signals. To test these predictions, we turned to real human EEG data to investigate 1) whether we could observe travelling waves in both IRF and EEG signals, as predicted by our model, and 2) whether their direction is task-dependent (e.g. visual processing vs. rest).

#### FW travelling waves during visual processing

Our group recently showed that alpha-band IRFs can be interpreted as FW travelling waves during visual perception (Lozano-Soldevilla and VanRullen, 2017). We first confirmed this result by applying our wave quantification method to a previously collected (Brüers and VanRullen, 2017) EEG dataset (INPUT) composed of 20 participants, who fixated a 6.25 second long white noise sequence of random luminance values for ~360 trials (fig.1A). For each electrode we considered the cross-correlation averaged over all trials. We obtained the 2D map and computed the log-ratio, as for the model, over seven midline electrodes (Oz to Fz – see fig.2A). Then, in order to estimate the proportion of reliable travelling waves, we computed the null distribution by shuffling the electrodes’ order 1000 times for each subject before re-computing the log-ratios. The difference between the real and the null distributions provides a statistical estimation of the proportion of reliable FW and BW wave events over and beyond what might be expected by chance, as captured by our surrogate distribution. In the INPUT dataset’s IRF, the comparison between the real and null distributions confirmed the presence of travelling waves (INPUT: Kolmogorov-Smirnov test, D=0.600, p<0.0008, fig.4B). Specifically, 48.6% of IRF data epochs showed evidence for FW waves over and beyond what could be expected by chance, and only 1.3% suggested BW waves. In addition, the predominance of FW waves was further confirmed by testing the distribution of mean log-ratios across participants against zero (BF=32.4, error=1.392e-4% -fig.2D).

**Figure 4.**
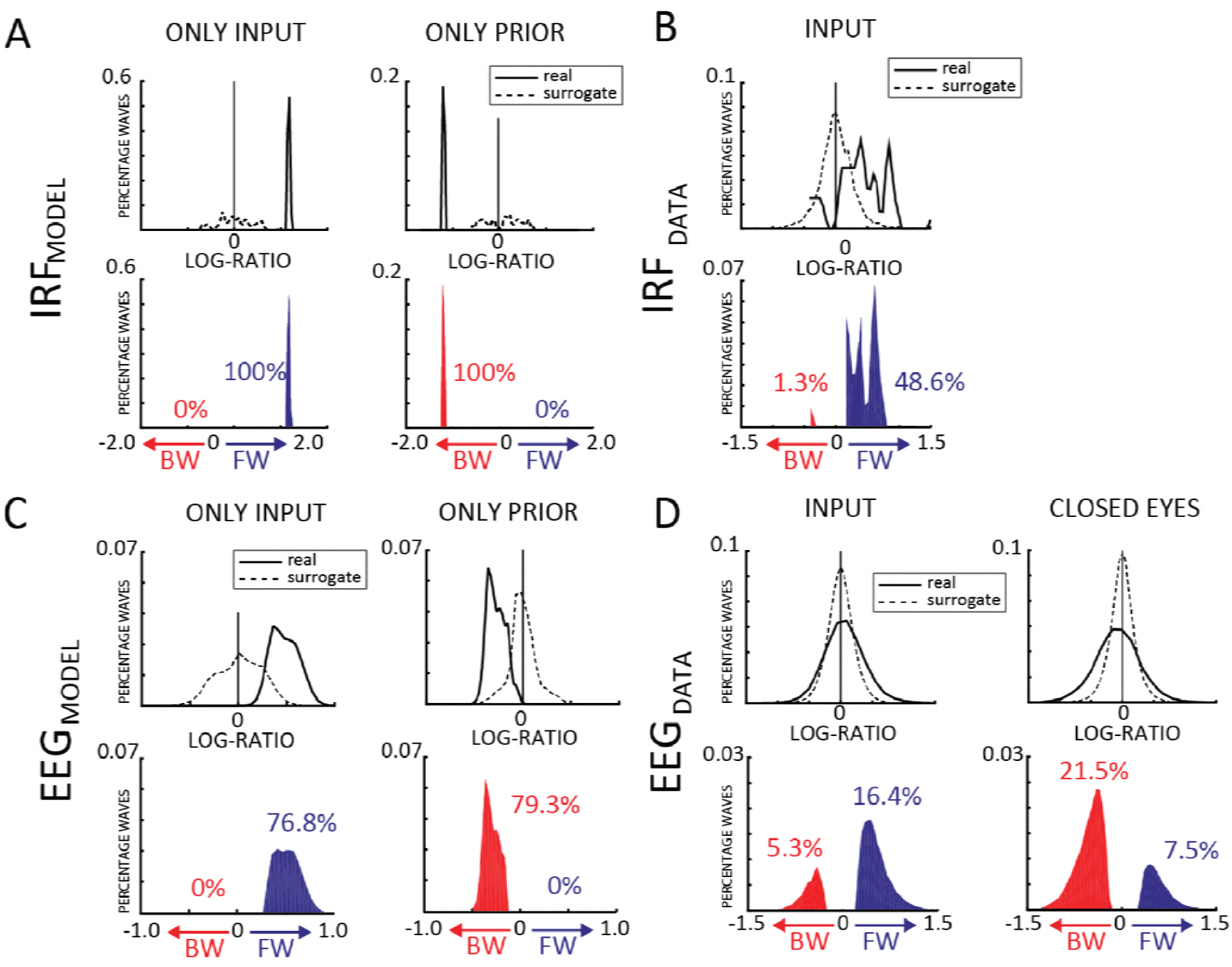
– Quantification of traveling waves in human EEG data and in model simulations. A) The first row shows the real (solid line) and shuffled (dashed line) distributions of log-ratios computed over the IRF obtained during 2 sets of simulations, cross-correlating prediction signals (a proxy for the EEG in our model) either with the sensory inputs (first column), or with the top-down prior signals (second column). The second row shows the difference between the two, focusing on positive values, i.e. the travelling wave events in the data that occur more often than predicted by the null distribution. The proportion of significant backward and forward waves are shown in red and blue, respectively. B) Same as A, but for the IRF computed in the INPUT human-EEG dataset (during human EEG experiments, we only have direct access to the visual input signals but the top-down priors, if any, remain unknown and cannot be used for cross-correlation). C,D) Same as A,B), but based on log-ratio values computed directly in 1s EEG epochs, rather than in the IRF (for the model, we use the prediction signals y_i_ as a proxy for the EEG signals). Experimental human EEG results (B,D) follow the same qualitative pattern as the simulations (A,C).

Once we confirmed the presence of FW travelling waves in the IRF measurements, we explored in the same dataset whether we could also demonstrate travelling waves in the raw EEG signals, as suggested by our model. Although several studies have reported travelling waves using intra-cortical recordings (Muller et al., 2018), there is so far little evidence from EEG studies (Alexander et al., 2013; Patten et al., 2012). Here, we applied a sliding window of 1 second (500ms overlap) to the continuous EEG data, and obtained approximately 4000 epochs per subject. For each epoch, we computed the log-ratio as for the IRF data, and again compared the real and null distributions. Remarkably, the Kolmogorov-Smirnov test used to assess a difference between the two distributions revealed the presence of travelling waves in the EEG data (INPUT: D=0.1584, p<0.0001; fig.4D). Particularly, 16.4% of the tested 1s EEG epochs showed reliable evidence in favor of FW waves, against only 5.3% of BW waves. The bias in favor of the FW propagation was confirmed by a Bayesian t-test between FW and BW waves (BF=134.3, error=1.055e-6%; see also fig.S2A). This result proves that direct EEG recordings can also reveal the presence of travelling waves, as suggested by the model’s simulations. Moreover, in line with the results using IRFs, the EEG data confirm that the bottom-up dynamics of perceptual processing are more strongly associated with FW than BW travelling waves.

#### BW travelling waves in resting-state EEG data

Our model simulations revealed that predominantly FW waves during stimulus processing are replaced by BW waves in the absence of stimulus. With the purpose of assessing the presence of BW waves in human EEG signals, we thus turned to a second dataset (CLOSED EYES) composed of 48 participants, who underwent a 1-minute recording with closed eyes. As for the previous dataset, we estimated the real and null log-ratios distributions after having computed the 2D maps for every 1 second time-window (500 ms overlap), over the same line of 7 central electrodes (Oz to Fz). Eventually we counted ~130 log-ratio values per subject. Notably, also in this dataset the comparison between the two distributions unveiled the presence of travelling waves (CLOSED-EYES: D=0.2149, p<0.0001; fig.4D) revealing, as predicted by our simulations, a preponderance of BW over FW waves (respectively 21.5% and 7.5% of the tested 1s EEG epochs), as confirmed by a Bayesian t-test between FW and BW waves (BF=74.2, error=2.632e-8%; see also fig.S2A). All in all, this result corroborates the predictions of our model regarding 1) the possibility to reveal travelling waves from raw EEG signals and 2) the presence of BW waves in the absence of sensory stimulation.

#### Human/Model comparisons

For the sake of comparison, we applied the same quantitative analyses to the modelling data as to the human EEG/IRF, obtaining qualitatively similar results. Notably, in the simulations we were able to compute an IRF both with the input and with the top-down prior signals (for human experiments, we only have direct access to the visual input signals, but the internal priors, if any, remain unknown). In all simulations, significant travelling waves were observed both in IRF data (D=0.9800, p<0.0001; fig.4A) as well as in EEG data (D=0.7982, p<0.0001; fig.4C). Similar to the CLOSED-EYES condition of the human EEG dataset, when only the top-down prior signal was present in the model, we observed exclusively BW waves: 79.3% of the tested 1s EEG epochs showed evidence for BW waves, and 100% of the IRF measurements (0% of EEG epochs or IRF measurements suggested FW waves). Conversely, when only the input was provided to the model, only FW waves were observed, in line with the results of the INPUT human EEG dataset: 76.8% of EEG epochs and 100% of IRF measurements revealed FW waves (with 0% of EEG epochs or IRF measurements supporting BW waves).

## Discussion

We presented a novel hypothesis suggesting that the ubiquitous alpha rhythm could reflect, in part, the computations involved in predictive coding. Long-lasting (~1s) alpha-band oscillatory reverberation of visual inputs, compatible with experimental observations (VanRullen and Macdonald, 2012), could be reproduced in a simple model with only short-term dynamics—each neuron only integrates information over ~20ms (neural time constant τ), and the delays for information transmission (ΔT) are also restricted to <20ms. A systematic exploration of parameter space revealed that IRF oscillations in the alpha (8-12Hz) frequency range systematically arise when τ and ΔT lie in their “biologically plausible” range. Therefore, we conjecture that it may simply not be possible for a biological brain, in which communication delays are non-negligible, to implement predictive coding, without also producing alpha-band reverberations. Moreover, a major characteristic of alpha-band oscillations, i.e. their propagation through cortex as a travelling wave, could also be explained by a hierarchical multi-layer version of our predictive coding model. The waves predominantly travelled forward during stimulus processing, and backward in the absence of inputs. These simulation results were remarkably matched by our human EEG data analyses (fig.5 provides a summary of the results), and are compatible with observations from other recent experimental studies (Halgren et al., 2017; Zhang et al., 2018).

**Figure 5.**
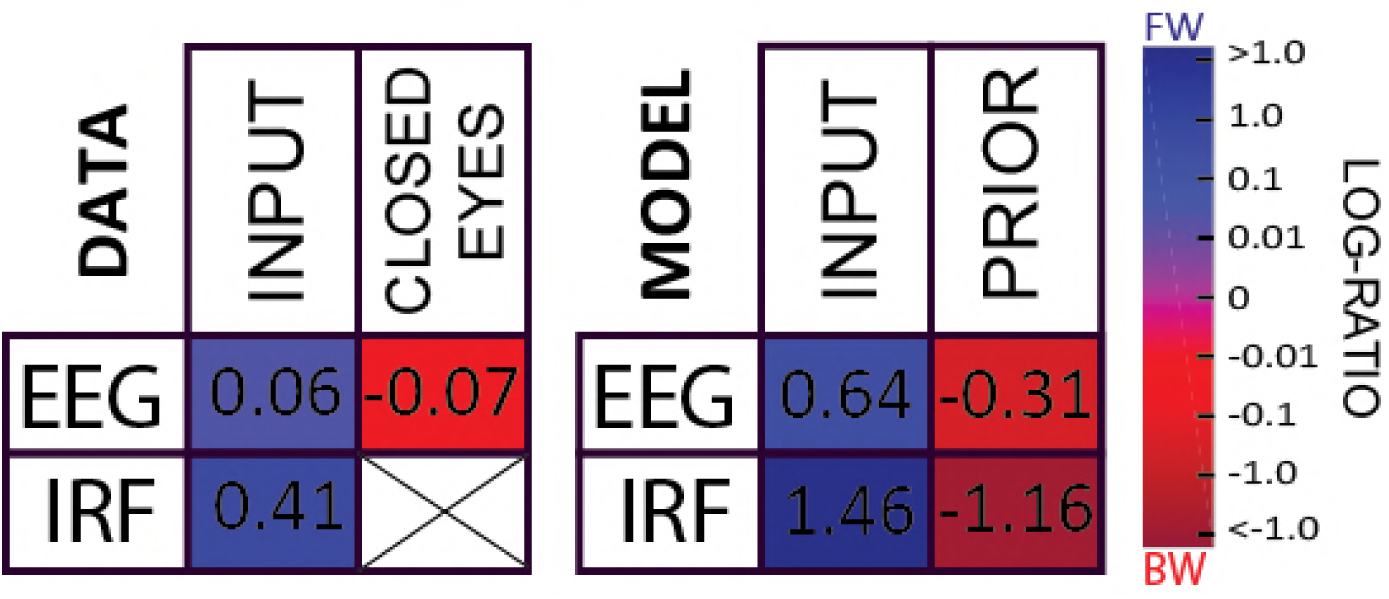
– Summary of the results. The tables summarize the results of the data (left) and simulation (right) for each dataset/condition. In each situation, the average log-ratio is presented numerically as well as color-coded. Red and blue squares indicate respectively a higher incidence of BW and FW waves. (As explained previously, it is not possible to compute an IRF in the CLOSED EYES condition).

### Generation of alpha (bidirectional interaction)

One of our simulations’ result refers to the generation of oscillations within a predictive coding framework enforced with temporal delays (Chalk et al., 2016). Specifically, our model posits that the bi-directional interaction between layers, influenced by communication delays, produces alpha-band rhythms. Electrophysiological studies have pointed at the origin of the alpha rhythms in the thalamic-cortical networks (Bollimunta et al., 2011; Steriade, 1997; Suffczynski et al., 2001), particularly involving the activity of the thalamic reticular nucleus (Lopes da Silva et al., 1980). The results of our 2-layers model (and the systematic exploration of its two free parameters τ and ΔT) may comply with this hypothesis, by considering respectively the lower and higher layer of the model as the thalamus and the primary visual cortex. However, other experimental evidence suggests that part of the alpha spectral power originates exclusively from cortical dynamics (Halgren et al., 2017; Lopes da Silva et al., 1980): in particular, pyramidal neurons in infra-granular and supra-granular layers of V2 and V4 have been implicated as alpha pacemakers (Bollimunta et al., 2011). Similarly, other cortical regions (such as V1 or S1) revealed the presence of alpha current generators, involved in both feedforward and feedback processes (Haegens et al., 2015, 2011; van Kerkoerle et al., 2014). The notion that alpha rhythms arise from the interaction of cortical layers is also in line with the results of our model, considering each of its layers as a different cortical region. Altogether, our model suggests predictive coding as a common computational framework involved in the generation of alpha oscillations, thus reconciling several apparently conflicting experimental findings on the origin of alpha.

### Types of travelling waves

Much work has been carried out regarding travelling waves, both in humans (mostly but not exclusively intra-cortical recordings – Muller et al., 2018) and in animals (e.g. turtles: Nenadic et al., 2003; Prechtl et al., 2000; cats: Roelfsema et al., 1997). Specifically, three different types of dynamic spatial propagation of brain signals have been characterized in the literature, and indiscriminately referred to as “travelling waves”. First, scalp ERPs propagating from posterior to anterior regions in response to stimulus onset have been interpreted as non-periodic travelling waves (Klimesch et al., 2007). Second, periodic sensory stimulations can produce oscillatory travelling waves that are confined to the same frequency as the entraining stimulus (Sato et al., 2012): even though they behave rhythmically, these waves are similar in principle to the first type above, as their oscillatory nature is not necessarily intrinsic, but directly related to the frequency characteristics of the generating stimulus. Lastly, truly periodic travelling waves have their own intrinsic frequency (e.g. within the alpha-band range, Lozano-Soldevilla and VanRullen, 2017), unrelated to the frequency content of the stimulus (if any). In this study, we investigated this last type of travelling waves, to understand the functional properties of intrinsically generated alpha-band oscillations.

### Generation of travelling waves

Ermentrout and Kleinfeld identified 3 different types of computational mechanisms that could lead to the generation of electrical travelling waves (Ermentrout and Kleinfeld, 2001). The first mechanism produces waves by virtue of a single neuronal oscillator, which spreads its rhythmic activity to neighboring and more distal regions with increasing time delays. In the brain, a potential candidate for such a role of pacemaker could be the above-mentioned cortical-thalamic circuit (Bollimunta et al., 2011; Destexhe et al., 1997; Jones, 2001; Muller and Destexhe, 2012). The second mechanism also suggests that the oscillatory wave originates as a result of a single oscillator, serially linked to other groups of neurons; however, whereas in the first model the wave’s motion was entirely due to varying communication delays from one oscillating region to all the others, in the second one the waves actually need to run through each region serially, with an increasing phase delay determined by the transmission delay between neighboring regions. Lastly, in the third mechanism each region is modeled as an independent oscillator with its own frequency and phase. In agreement with the Kuramoto model (Kuramoto, 1984), in this scenario the wave propagation follows the phase shifts between each region, which in turn depends on their relative frequencies, travelling from higher to lower frequencies. Remarkably, despite its numerous assumptions (i.e. weak coupling between regions, nearly identical oscillatory properties across regions), this last model (Kuramoto, 1984) has been successfully applied in several computational brain studies (Breakspear et al., 2010; Cumin and Unsworth, 2007). Although our model does not exactly fit within any of these three categories, it does share some similarities with the last two. On the one hand, the local interactions between neighboring regions with a constant communication delay relates to the second approach; on the other hand, the movement of the travelling wave arises naturally as in the last model from the communication between separately oscillating layers. Importantly though, in our model none of the units is an oscillator or pacemaker per se; oscillations arise from bidirectional communication with temporal delays. This property marks a decisive difference with all three previous models. Lastly, in our implementation the wave’s direction does not depend on a gradient of time delays or a gradient of frequencies; instead, the same model can produce both FW or BW waves depending on the signals provided: FW waves originate from perceptual inputs, BW waves from top-down activity.

### Functional role of travelling waves

The travelling direction of the waves appears important in determining their functional role. In a recent study (Halgren et al., 2017), intra-cortical recordings from epilepsy patients during quiet wakefulness revealed alpha-band travelling waves propagating “backward” from higher-order antero-superior cortex to lower-order occipital poles. Analyzing recordings simultaneously from cortex and the pulvinar, Halgren and colleagues concluded that alpha-band travelling waves 1) originate in the cortex, and 2) reflect feedback processing between cortical regions. However, another recent human intracranial study from Zhang and colleagues (Zhang et al., 2018) reported theta-alpha travelling waves (2 to 15 Hz) that could propagate “forward”, from posterior to anterior brain regions during the cue period of a visual memory task. Although these two studies appear contradictory in terms of the propagation direction, our model can easily reconcile these findings, by positing that alpha-band travelling waves emerge as a result of layer-by-layer interactions in a hierarchical system, and their direction is related to their functional role: forward to convey input signals (and the prediction residuals) during sensory processing and task engagement; backward to convey top-down priors and expectation signals, most visible during rest and in the absence of inputs.

### Speed of travelling waves

The speed of cortical travelling waves appears to depend on several factors, as reflected by the large range of values reported in the literature (from 0.1 to 10m/s). One important distinction regards whether the travelling waves are recorded at a macroscopic (i.e. whole-brain) or mesoscopic level (i.e. within single regions in the cortex) (Muller et al., 2018). In the first case, M-EEG or ECoG recordings allow to determine their propagation, which supposedly occurs through myelinated axons of white matter fibers: consequently, their speed spans between 1 to 10 m/s (Muller et al., 2018; Patten et al., 2012). Conversely, techniques with higher spatial resolution, such as voltage sensitive dye imaging (Shoham et al., 1999) or multi-electrode arrays (Borroni et al., 1991), grant access to local, slower travelling waves, whose speed ranges from 0.1 to 0.8 m/s (Muller and Destexhe, 2012; Zhang et al., 2018), in agreement with the axonal conduction speed of unmyelinated long-range and horizontal fibers within the cortex (Girard et al., 2001). In addition, another important factor potentially affecting the measured travelling waves speed is cortical source mixing, which is especially pronounced when recording from the scalp (i.e. M-EEG): as every recording channel picks up not only local but also more distal cortical signals, we should expect an apparently faster speed at the scalp level than the true underlying speed of the travelling wave over the cortex. This could explain why the waves seem to traverse the 7 layers of our model much more slowly than they go through our 7 scalp-level electrodes in the human EEG recordings (compare Figure 2A to Figure 3B). In support of this view, we simulated EEG source mixing in our model to assess the corresponding changes in wave speed, and then compared the results with real EEG data (see Supplemental Figure S3). By assuming a layer-to-layer distance of 2cm, roughly equivalent to the distance between neighboring cortical regions, our model produces travelling waves whose speed falls in the mesoscopic range (~0.6 m/s, see fig.S3), comparable with experimental cortical recordings (Muller and Destexhe, 2012; Zhang et al., 2018). However, after remapping the model deep layers to superficial electrodes by means of a weighted average (thus simulating cortical source mixing, fig.S3, panel B), the speed of the traveling waves increased, now falling within the range of macroscopic waves (~2.2 m/s). Importantly, this linear averaging, despite increasing the apparent speed, did not significantly affect the waves’ direction or their log-ratios. Furthermore, the speed of the scalp-level travelling waves as simulated by our model was now comparable with the one observed in our experimental human EEG dataset (~2.2 m/s, see fig.S3, panel D).

### Relation to other predictive coding models

Due to its streamlined architecture, our predictive coding model produces a single oscillation whose frequency depends on the chosen parameters: typically in the alpha band for biologically plausible values (Figure 1D). The same alpha oscillation thus carries top-down predictions down the hierarchy, and bottom-up prediction residuals up the hierarchy, resulting in two alpha travelling waves moving in opposite directions. As explained above, this is compatible with recent experimental reports of both feedforward and backward alpha travelling waves in human intracranial studies (Halgren et al., 2017; Zhang et al., 2018). However, there is also a growing number of studies reporting that faster gamma oscillations (~30-80Hz) are specifically involved in feed-forward signal transmission, while alpha-and beta-band (13-30Hz) rhythms convey top-down information (Bastos et al., 2012; Buffalo et al., 2011; Michalareas et al., 2016; van Kerkoerle et al., 2014). These dynamics have been appropriately captured by a more detailed predictive coding model, the “canonical cortical microcircuit” model (Bastos et al., 2012; Friston, 2005), which includes different types of excitatory and inhibitory neurons, as well as detailed laminar circuitry in each brain region. An important next step would thus be to explore the existence, frequency and direction of travelling waves in a hierarchical version of the canonical microcircuit model. One can speculate that such a model could account for both the bi-directional nature of alpha travelling waves across the hierarchy (as observed in our model and in the above-cited human intracranial studies), and for the prevalence of gamma-band signals in the feed-forward communication between sending and receiving layers of two consecutive brain regions (as measured e.g. using Granger causality; Bastos et al., 2012; Michalareas et al., 2016). Finally, future versions of our model could also expand the number of simulated neurons in each layer, together with a retinotopic organization and spatially selective receptive fields (as in Rao and Ballard, 1999), in order to process spatially as well as temporally structured inputs. This should provide a more complete understanding of predictive coding and oscillatory travelling waves in relation to essential visual functions such as object recognition or categorization.

## Materials and methods

### Model and simulations

Predictive coding (Rao and Ballard, 1999) postulates a hierarchical architecture in which higher layers predict the activity of lower layers, and the residuals (i.e the difference between the prediction and the actual activity) are carried over to update the next prediction. In our model, illustrated in fig 1B and 3A, the residual is defined as:

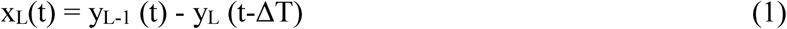

where L indexes the layers, and ΔT represents the temporal communication delay between them. For simplicity, the delay is assumed here to be symmetric in both forward and backward directions, but it can easily be shown that the oscillatory behavior chiefly depends on the sum of forward and backward delays (e.g. comparable oscillatory dynamics would be found for symmetric delays with ΔT=12ms, or for asymmetric delays with ΔT_forward_=16ms and ΔT_backward_=8ms). For the consistency of notations, we pose y_L-1_(t) = INPUT(t) when L=1. The prediction y_L_, as shown in equation 2, is updated based on the bottom-up residual x_L_ (with a delay ΔT), and on the difference between its prediction and the prediction from the next higher layer, which can be considered as a top-down prior:

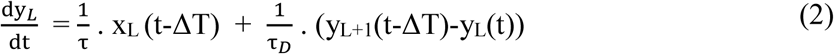

At the last layer, the prior y_L+1_ can serve to represent a generative endogenous process, arising from higher-level brain regions that are not part of the model; in our simulations, this top-level prior could be imposed as a time-varying signal or set to 0 (see fig. 3A). In order to facilitate measurements of cross-correlations (IRF) with this prior, and comparison with the stimulus-induced IRF, when different than 0 we set the time-varying top-level prior to a white-noise signal with statistics similar to (but independent from) those of the input signals.

Besides ΔT, two other parameters play a crucial role in the model: τ and τ_D_ which describe the temporal dynamics (time constants) of neuronal integration and decay, weighting respectively the residual computed from the lower-layer, and the prediction from the higher layer. Note that when the prior y_L+1_ is set to 0, the second term in Equation (2) acts as a decay term, which would ensure that the prediction y_L_(t) returns to zero in the absence of inputs. For this reason, and to take into account the fact that higher-level brain signals typically vary slower than low-level input signals (Gauthier et al., 2012; Kiebel et al., 2008; McKeeff et al., 2007; Murray et al., 2014), we set the time constant τ_D_ to a fixed value of 200ms. In all simulations, equations (1) and (2) were solved numerically with a 1ms time step.

The results of the parameter search in the 2-layers model (fig.1D) were obtained presenting, for each pair of τ and ΔT, 200 different white noise luminance sequences of 3s. The model’s IRF was computed by cross-correlating each luminance sequence with layer 2’s output y(t), and averaging the results over the 200 trials. Regarding the results of the multilayer model (fig.4 A,C), we investigated 2 possible scenarios, in which either an input or a prior signal was presented to the model for 6s on each of 200 simulated trials. Consequently, we computed the IRF by cross-correlating each layer’s output y_L_(t) with one of the two signals. We then defined 2D-maps (fig.2A-B) stacking up the temporal signals from all 7 model layers. We thus obtained two 2D-maps, one for each scenario, i.e. IRF with input or IRF with prior. Moreover, we computed 2 additional 2D-maps considering the raw (simulated) EEG signals (no cross-correlation) in place of the IRF, separately for each scenario (in this case, the temporal signals were obtained with a 1s sliding window with 500ms overlap, yielding 11 distinct analysis windows for each trial). For each of these maps we then computed a log-ratio that quantifies the presence and the direction of the waves (see below). Log-ratio distributions were obtained pooling together the results over the 200 simulated trials. We followed the same procedure for the null distribution, obtained by randomly shuffling the order of the layers before computing the log-ratios.

### Analytical solution of the (simplified) model

An analytical solution for our system of delayed differential equations (1) and (2) exists (Kaplan and Yorke, 1974) for a 2-layer model under two simplifying assumptions: 1) we neglect the second term of equation (2), given that τ_D_ >> τ; 2) the integral over time of the input signal amounts to 0 (e.g. white noise centered on 0). This leads to the following equation, obtained by merging the simplified equations (1) and (2):

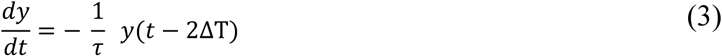

If we consider as a general solution the exponential function in (4a), we can compute each side (respectively 4b and 4c) of the equation in (3) as:

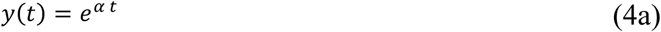

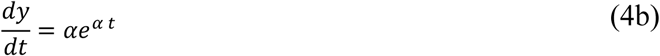

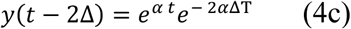

By replacing (4b) and (4c) in (3), we obtain equation (5a), simplified in (5b):

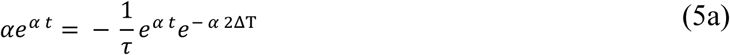

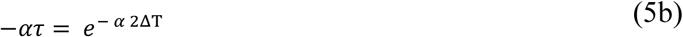

A “pure” oscillatory solution would correspond to a situation where α is an imaginary complex number, that is, α = 0 + iω (where ω is the oscillatory frequency). Given (5b), this leads to:

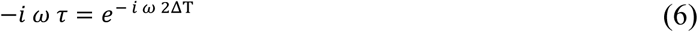

Eventually, applying Euler’s formula, we derive two equations by separately considering the real (7a) and the imaginary (8a) parts. Since the real part is equal to 0, we obtain

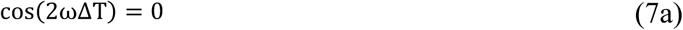

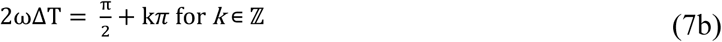

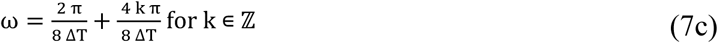

whose first oscillatory solution (with lowest frequency) is:

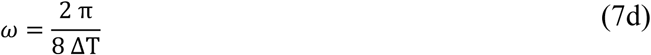

In plain English: if the model solution is oscillatory, it will most likely oscillate with a period around 8ΔT.

Regarding the imaginary part, we can write:

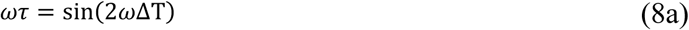

which, given Equation (7a), must equal 1 or −1. As both ω and τ are positive, we conclude that:

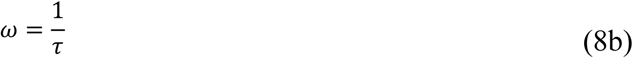

Combining (7d) with (8b) provides equations (9a-b)

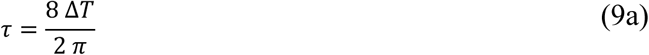

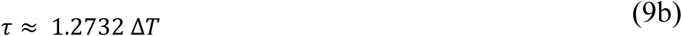

In conclusion, oscillatory solutions exist in a region of parameter space (ΔT, τ) slightly above the diagonal (τ=ΔT). The period of the oscillation is 8ΔT, which will lie in the alpha-band (8-13Hz) whenever ΔT is between 10 and 15ms. In that case, Equation (9b) suggests that τ should be around 13-20ms. Therefore, the equations theoretically confirm the results of the numerical parameter search (fig. 1D), defining the region of parameter space where solutions oscillate in the alpha band.

### Experimental datasets

#### Participants

The INPUT dataset was composed of EEG recordings from 20 volunteers (aged 23–39 years old with a mean age of 28, 10 women, 5 left handed), the CLOSED-EYES dataset from 48 volunteers (aged 20– 43 years old with a mean age of 27.8, 25 women, 7 left handed). All subjects reported normal or corrected to normal vision and no history of epileptic seizures or photosensitivity. All participants gave written informed consent before starting the experiment, in accordance with the Declaration of Helsinki. This study was carried out in accordance with the guidelines for research at the “Centre de Recherche Cerveau et Cognition” and the protocol was approved by the committee “Comité de protection des Personnes Sud Méditerranée 1” (ethics approval number N° 2016-A01937-44).

#### Stimuli and protocols

Results from the INPUT dataset have been published elsewhere (Brüers and VanRullen, 2018, 2017), whereas the second, CLOSED-EYES dataset presents a mixture of published and unpublished data. Regarding the INPUT dataset, the experiment was composed of two sessions of 8 experimental blocks of 48 trials each, having a total duration of about 1 hour. Participants viewed 6.25 second long random (white-noise) luminance sequences presented on a cathode ray monitor, positioned ~57 cm from the subject, with a refresh rate of 160 Hz and a resolution of 640 × 480 pixels. Each sequence had a flat power spectrum up to 80 Hz and was designed using MATLAB custom scripts and displayed using the Psychophysics Toolbox (Brainard, 1997). The stimuli were presented on a black background in a peripheral disk with a diameter of 7°, whose center was at 7° of eccentricity above the fixation point. Concerning the CLOSED-EYES dataset, participants were asked to close their eyes for 1 minute while recording their spontaneous brain activity (this is done routinely in our lab to measure each participant’s individual alpha frequency). After the recording ended, participants performed other tasks unreported in this manuscript.

#### EEG recording and Pre-processing

In both datasets, brain activity was recorded using a 64 channels active BioSemi electro-encephalography (EEG) system (1,024 Hz digitizing rate, 3 additional ocular electrodes). The following pre-processing steps were applied to all subjects of the INPUT dataset using the EEGlab toolbox (Delorme and Makeig, 2004) in Matlab. Once the noisy channels had been rejected and interpolated (when necessary), the data was offline down-sampled to 160 Hz to match the presentation rate of luminance stimuli and thus facilitate the cross-correlation of the two signals. A notch filter [47Hz −53Hz] was then applied to remove power line artifacts. We applied an average-referencing and removed slow drifts by applying a high-pass filter (>1 Hz). In the first dataset (i.e. INPUT), data epochs were created around each white-noise sequence (from −0.25 to 6.5 s) and the baseline activity was subtracted (i.e., mean activity from −0.25 s to 0 before trial onset). Finally, the data was screened manually for eye movements, blinks and muscular artifacts and whole epochs were rejected as needed: on average 20/384 trials were rejected per subject. For the second dataset (CLOSED-EYES), preprocessing was limited to noisy channels rejection, and a similar filtering process as in the previous dataset: after a notch filter [47Hz −53Hz] was applied, a high-pass filter (>1Hz) removed slow drifts.

#### Log-ratio

We computed log-ratios of each 2D map (fig.2A-B) in order to assess and quantify the presence of travelling waves in both EEG signals and simulations. In order to create 2D maps of the human EEG data we considered either the raw EEG signals from 7 midline electrodes (posterior to frontal: Oz, POz, Pz, CPz, Cz, FCz, Fz), or their cross-correlation with the input sequence (i.e. IRFs). Regarding the simulations, we utilized either the prediction signals from each layer of the model or their cross-correlation (IRF) with either the input or the prior signals. Each 2D map for the raw EEG (or its model equivalent) was computed with a sliding window of 1 second and an overlap of 500ms (x-axis). 2D maps for IRFs were obtained directly by stacking the 7 IRF time courses (from 7 EEG electrodes or from 7 model layers), as the support window of the IRF was only 1-s long. The maximum value of the upper-left quadrant of the 2D-FFT was extracted to represent the amount of feedforward (FW) signal propagation across layers/electrodes, whereas the maximum value of the lower-left quadrant revealed the amount of backward (BW) waves. Finally, we computed the log-ratios by dividing the FW maximum value by the BW maximum value, and taking the log of the result: positive and negative ratios revealed respectively FW and BW waves.

## Acknowledgments

This work was funded by an ERC Consolidator Grant P-CYCLES number 614244. We are grateful to Sasskia Brüers for EEG data collection, and to Andreas Herz for pointing out the existence of an analytical solution to the delayed differential equations, and for helping derive it. Parts of this work were done while RV was visiting the Simons Institute for the Theory of Computing.

## Supplementary figures

**Figure S1.**
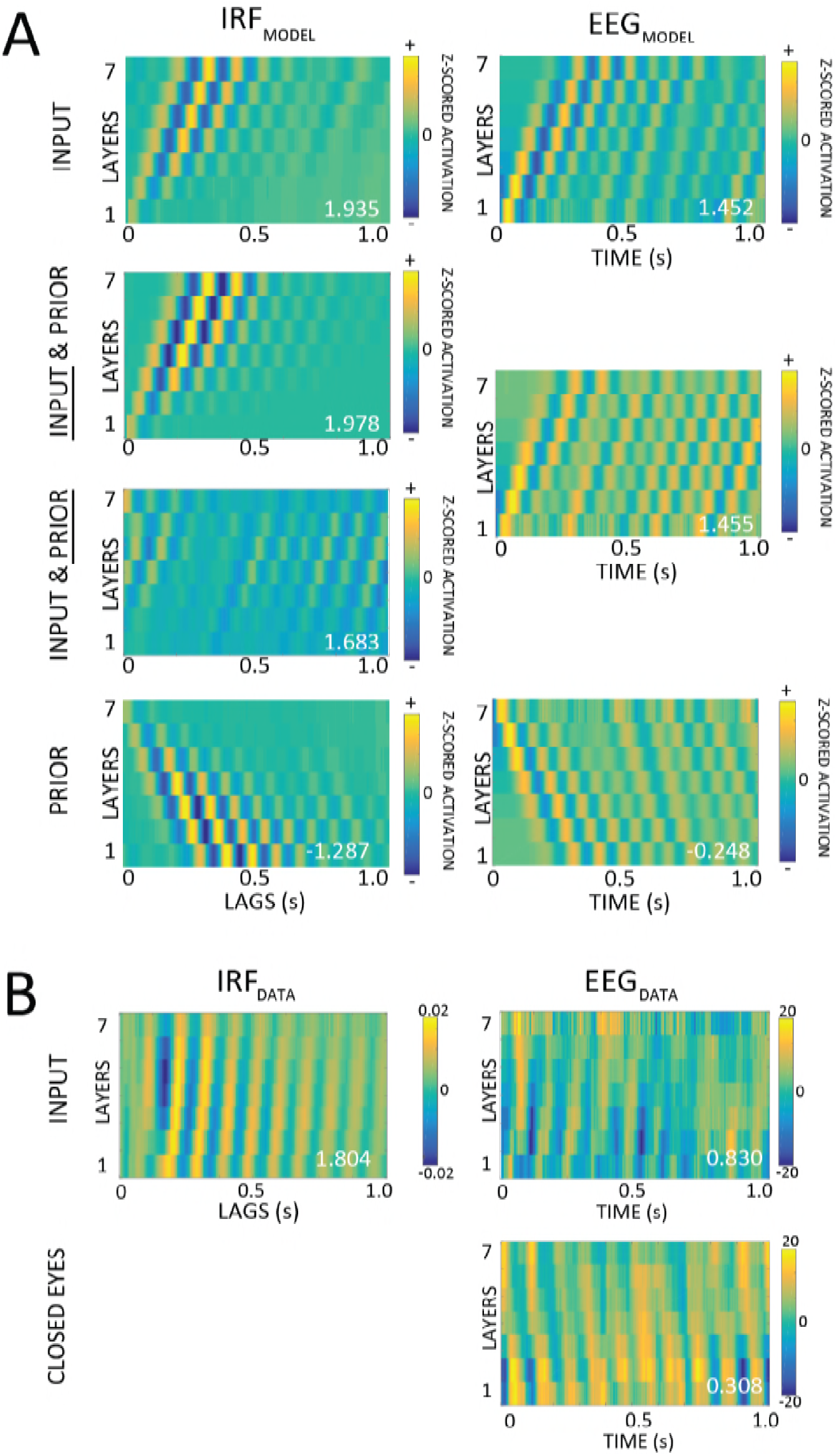
– Overview of 2D maps from all the simulations and experimental data. A) The picture reports all the possible simulation scenarios. The left column shows the 2D maps of the IRF, whereas the right one the 2D maps of the EEG (i.e. the prediction signals from each model layer). Each row shows the results of a different simulation, with either the input signal (first row), the top-down prior signal (last row), or both together (middle panels; although there is only one EEG signal in this situation, it can be cross-correlated with either the input or the prior signals, hence the two panels in the left column). B) The first row shows IRF (left) and EEG (right) 2D maps from one representative participant in the INPUT dataset. The second row shows another representative participant in the CLOSED-EYES dataset; only the EEG map is displayed, since we do not have access to either external input signals or internal top-down priors to cross-correlate with the EEG and derive an IRF.

**Figure S2.**
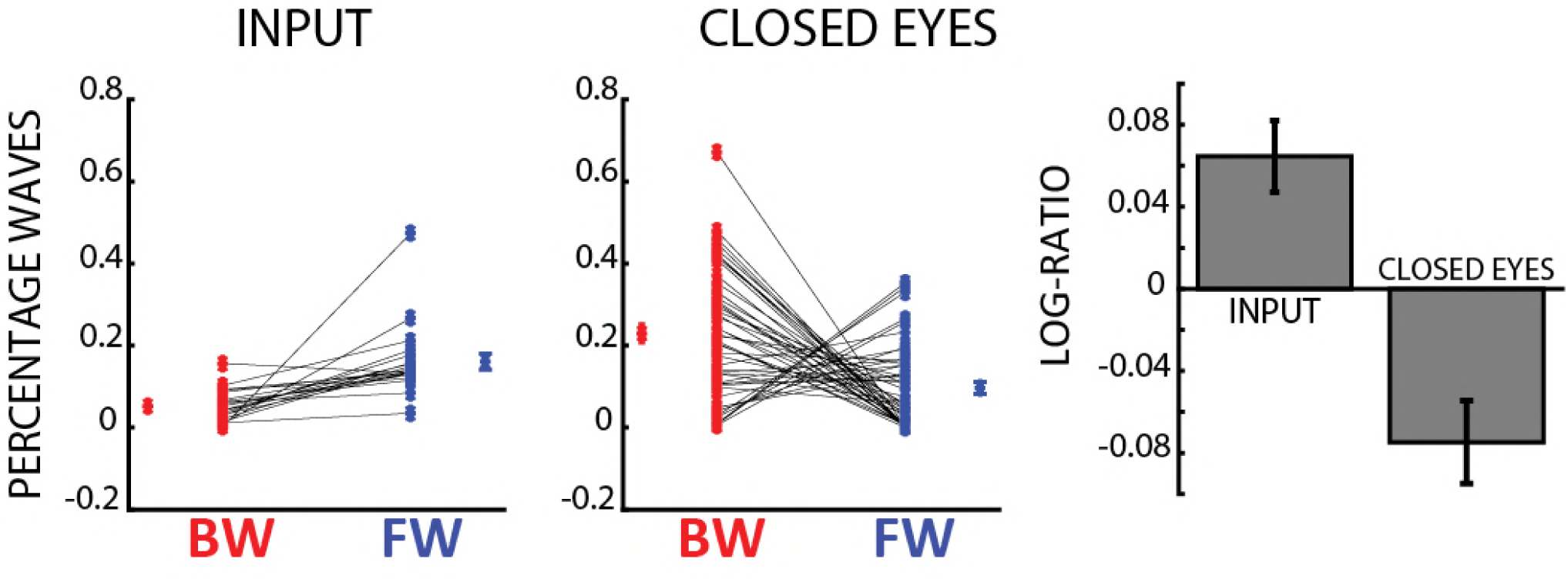
– Direction of traveling waves in human EEG data. The first 2 panels show the proportion of backward (BW, in red) and forward (FW, in blue) travelling waves respectively in the INPUT and CLOSED-EYES dataset for each participant, as computed in figure 4D. Each value represents the percentage of waves that occur above chance level, when compared to the null distribution (see figure 4 and Material and Methods for details). In the INPUT dataset most participants have a larger number of FW waves than BW, whereas the CLOSED-EYES dataset reveals an opposite trend. The mean +/− SE log-ratios of both datasets result corroborated this result, as shown in the right panel.

**Figure S3.**
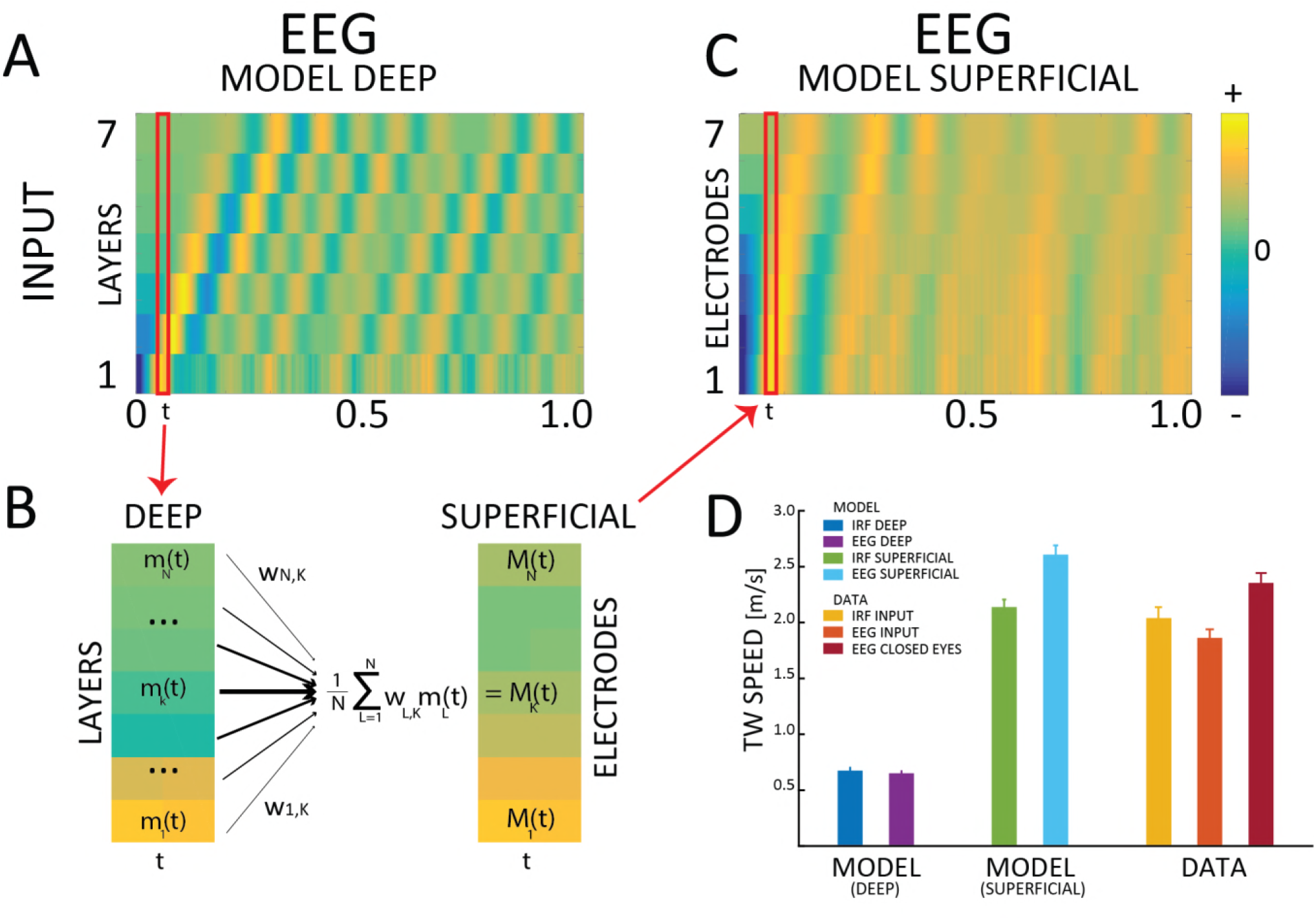
– Speed of travelling waves. A) Instead of assuming that distinct model layers correspond to distinct scalp-level electrodes, each layer of the model can be interpreted as a cortical brain region, and consequently each prediction sequence as a cortical recording. Under these assumptions, the speed of the travelling wave, computed assuming a distance of ~2cm between neighboring cortical regions, is ~0.6m/s (model EEG in the picture, but similar results are obtained with IRF data). B) Applying a moving linear weighted average for every layer at each time point (the red box highlights an example) determines a transformation in which each new layer (i.e. electrode) is influenced by neighboring layers, in the same way as superficial EEG electrodes are presumably affected by multiple deep cortical sources. In this transformation, the contribution of each layer is weighted based on its distance to the electrode. The contribution of the kth layer for the kth electrode is equal to 1, and it decreases progressively with distance to 0.8, 0.6 and 0.4 (respectively for layers k±1, k±2 and k±3). The contribution of layers further away was set to 0. C) The transformed map reveals a travelling wave whose speed is faster, and more compatible with EEG recordings. Importantly, the wave direction and the log ratios are not significantly influenced by such transformation. D) The travelling wave speed of the model before (blue and violet) and after (green and cyan) the transformation, compared to the speed of all the significant waves in our experimental datasets (i.e. epochs whose log-ratio was above chance level, as estimated by the surrogate distributions). We estimated an average electrode-to-electrode distance of ~4cm for the real data and the superficial model simulations, and a layer-to-layer distance of ~2cm for the deep model.

